# S100A8/A9 Inhibition Reduces Splenic Myelopoiesis and Improves Outcomes After Stroke

**DOI:** 10.64898/2025.12.03.692236

**Authors:** Hyun Ah Kim, Annas Al-sharea, Hannah X. Chu, Sung-Chun Tang, Samoda A. Rupasinghe, Shenpeng R. Zhang, Prabhakara R Nagareddy, Grant R. Drummond, Thiruma V. Arumugam, Andrew J. Murphy, Christopher G. Sobey, Man K.S. Lee

**Affiliations:** Centre for Cardiovascular Biology and Disease Research, La Trobe Institute for Molecular Sciences, La Trobe University, Bundoora, VIC, Australia; Department of Microbiology, Anatomy, Physiology and Pharmacology, School of Agriculture, Biomedicine and Environment, La Trobe University, Bundoora, VIC, Australia; Haematopoiesis and Leukocyte Biology, Baker Heart and Diabetes Institute, Melbourne, VIC, Australia; Stroke Center, Department of Neurology, National Taiwan University Hospital, Taipei, Taiwan; Department of Internal Medicine, Section of Cardiovascular Diseases, University of Oklahoma Health Sciences Center (OUHSC), Oklahoma City, OK, USA; Baker Department of Cardiometabolic Health, The University of Melbourne, Melbourne, VIC, Australia; Department of Diabetes, Monash University, Melbourne, VIC, Australia; Baker Department of Cardiovascular Research, Translation and Implementation, La Trobe University, Bundoora, VIC, Australia

**Keywords:** S100A8/A9, ABR-215757, paquinimod, inflammation, stroke, middle cerebral artery occlusion

## Abstract

**Background:** Neutrophils are among the earliest immune cells to infiltrate the ischemic brain and contribute to secondary neuronal damage. The alarmin S100 calcium-binding protein A8/A9 (S100A8/A9), predominantly released by neutrophils, is upregulated during this process. Although the bone marrow is recognised as the principal site of neutrophil production via myelopoiesis, the role of the spleen as an immune-responsive organ remains incompletely understood.

**Methods:** In this study, we employed a transient middle cerebral artery occlusion (MCAO) model in male C57Bl/6 mice and examined immune responses 24 hours post-stroke in the blood, bone marrow and spleen using flow cytometry. To understand the role of S100A8/A9 in modulating stroke-induced myelopoiesis, we administered a small molecule inhibitor of S100A8/A9, ABR-215757, before and after stroke.

**Results:** Neutrophils and S100A8/A9 were found in the infarcted brain tissue. Interestingly, we observed a marked increase in splenic neutrophils, accompanied by an expansion of myeloid progenitors, indicating activation of extramedullary myelopoiesis. Given our previous work showing that S100A8/A9 promotes myelopoiesis, we pharmacologically inhibited S100A8/A9 to determine if this would modulate stroke-induced myelopoiesis. Treatment with ABR-215757 at 24 hours post-stroke led to reduced splenic myelopoiesis, reversed neutrophilia, enhanced forelimb grip strength, and a one-third reduction in infarct size.

**Conclusion:** These findings identify the spleen as a key contributor to neutrophil production following stroke and suggest that targeting S100A8/A9 may attenuate post-stroke inflammation and improve neurological recovery.

**Highlights:**

- Stroke induces extramedullary myelopoiesis in the spleen, not femoral bone marrow.
- Neutrophil-derived S100A8/A9 drives splenic myelopoiesis after ischemic stroke.
- Pharmacological blockade of S100A8/A9 with ABR-215757 reduces neutrophilia.
- Inhibition of S100A8/A9 lessens infarct size and improves neurological recovery.
- Human stroke tissue confirms S100A8/A9 accumulation with neutrophil infiltration.

## Background

Ischemic stroke, a leading cause of mortality and long-term disability, is caused by a sudden disruption of cerebral blood flow that leads to neuronal injury and a complex inflammatory response. While restoring blood flow via thrombolysis (tPA)^1^, anti-platelet drugs^2^, or thrombectomy^3^ remain the primary therapeutic approaches, these strategies do not address the secondary inflammation-driven brain injury that worsens stroke outcomes.

Neutrophils and monocytes infiltrate the ischemic brain within the first few hours of stroke onset^4^, exacerbating injury through disruption of the blood-brain barrier (BBB), cerebral edema, and neurotoxic cytokine release^5^. This pathological mechanism involves factors released by neutrophils, including reactive oxygen species (ROS) (superoxide), proteases (matrix metalloproteinases, elastase), cytokines (interleukin-1β (IL-1β), IL-6, IL-8, tumour necrosis factor-alpha (TNF-α)) and chemokines (CCL2, CCL3, CCL5)^6^. Clinically, elevated neutrophil counts correlate with stroke severity^7^, infarct size^8^, and worse functional outcomes^9^. Despite their role in driving inflammation, the source of these myeloid cells and the mechanisms regulating their production remain poorly understood.

Emergency myelopoiesis, the rapid expansion of myeloid progenitors in response to acute inflammation, is a well-characterised phenomenon in several diseases including infection^10^ and myocardial infarction^11,12^. Traditionally, the femoral bone marrow was considered the primary site of post-stroke neutrophil and monocyte production. However, recent studies from the Nahrendorf group challenge this view, demonstrating that the skull bone marrow, instead of the femoral bone marrow in mice, serves as the dominant source of neutrophils entering the infarct during the first 24 hours^13^. These findings suggest that post-stroke myelopoiesis is compartmentalised, potentially involving multiple distinct hematopoietic reservoirs.

Beyond the bone marrow, the spleen has also emerged as a key immune organ in stroke pathology^14^. The marked ability of the spleen to contract in both preclinical models and in stroke patients has been shown to result in the release of stored leukocytes into the circulation, exacerbating inflammation. However, whether the spleen acts as a reservoir or as an active site of myeloid cell production remains unknown.

S100 calcium-binding protein S100A8 and S100A9 are damage-associated molecular pattern (DAMP) molecules, that are present in increased levels in several inflammatory and autoimmune states^15^. S100A8 and S100A9 constitute approximately 40% of the cytosolic protein content in human blood neutrophils and 1% in monocytes^16^. Once released extracellularly, S100A8/A9 functions as an endogenous agonist to bind to toll-like receptor 4 (TLR4)^17^ and the receptor for advanced glycation end products (RAGE)^18,19^, driving emergency myelopoiesis as seen in infection^10^ and cardiovascular disease^11^. In stroke, clinical studies have demonstrated elevated plasma S100A8/A9 levels that correlate with disease severity^20^. While we and others have previously demonstrated that S100A8/A9 promotes emergency myelopoiesis in the bone marrow^12,21,22^, its role in stroke remains unexplored.

Given the ability of S100A8/A9 to regulate myeloid expansion, we hypothesised that S100A8/A9 may act as a key signal driving neutrophil and monocyte production in response to stroke. Herein, we first assessed the expression of S100A8/A9 and neutrophils in human and mouse brains after ischemic stroke and their role in ischemic brain injury. We then explored stroke-induced myelopoiesis and the impact of S100A8/A9 on the formation of neutrophils and stroke outcome.

## Results

### Stroke increases S100A8/A9 expression in mouse and human ischemic brain

Transient middle cerebral artery occlusion (tMCAO) in mice resulted in robust upregulation of S100a8 and S100a9 mRNA in the ischemic hemisphere at 24 h compared with the contralateral side (Figure 1A). Immunohistochemistry confirmed accumulation of S100A8/A9 together with infiltrating neutrophils in the infarct region (Figure 1B). Similar findings were observed in post-mortem human stroke tissue, where S100A8/A9 was detected alongside neutrophil infiltration (Figure 1C), supporting the translational relevance of this pathway.

**Figure 1.**
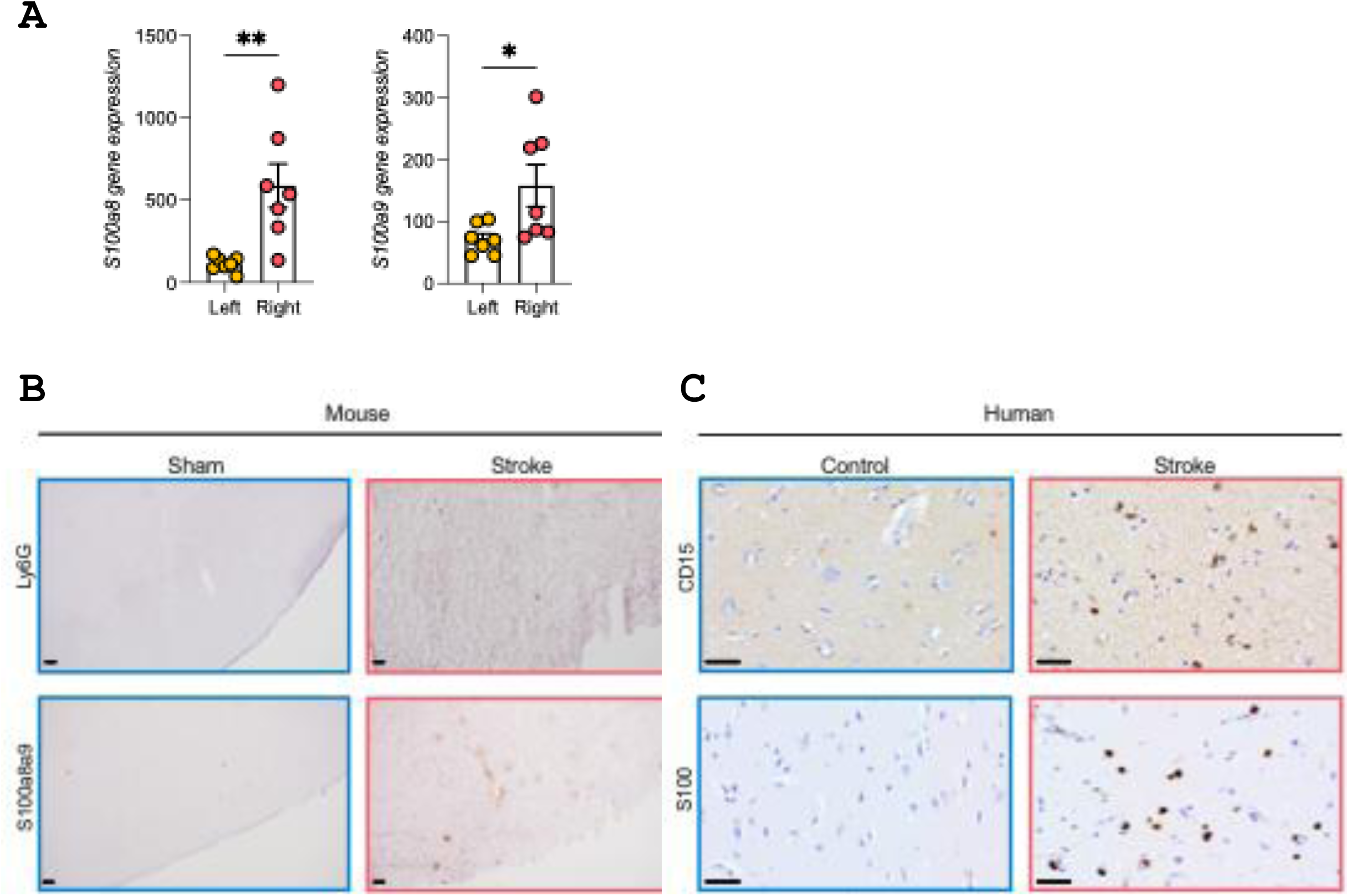
S100A8/A9 is upregulated in mouse and human ischemic brain. (A) *S100a8/a9* mRNA expression in ipsilateral (right) vs contralateral (left) hemispheres 24 h after tMCAO. (B) Mouse immunohistochemistry shows the expression of S100A8/A9 and neutrophils (Ly6G+) in infarcted cortex. (C) Human post-mortem stroke brain also shows the expression of S100A8/A9 and neutrophil (CD15) in the infarct. Student’s t-test. *p<0.05, **p<0.01. n=6-8 per group. Scale bar = 50µm.

### Stroke induces splenic, but not bone marrow, myelopoiesis

Stroke was associated with a marked increase in circulating neutrophils, but no change in monocytes at 24 h (Figure 2A). Flow cytometry revealed expansion of hematopoietic stem and progenitor cells (LSKs) in femoral bone marrow (Supplementary Figure 1A); however, downstream myeloid progenitors and mature neutrophils were not significantly altered (Supplementary Figures 1A and 1B), consistent with previous reports. In contrast, the spleen exhibited a striking expansion of myeloid progenitors (CMPs, GMPs; Figure 2B) as well as increased monocytes and neutrophils (Figure 2C), indicating that splenic myelopoiesis is a key contributor to systemic neutrophilia after stroke.

**Figure 2.**
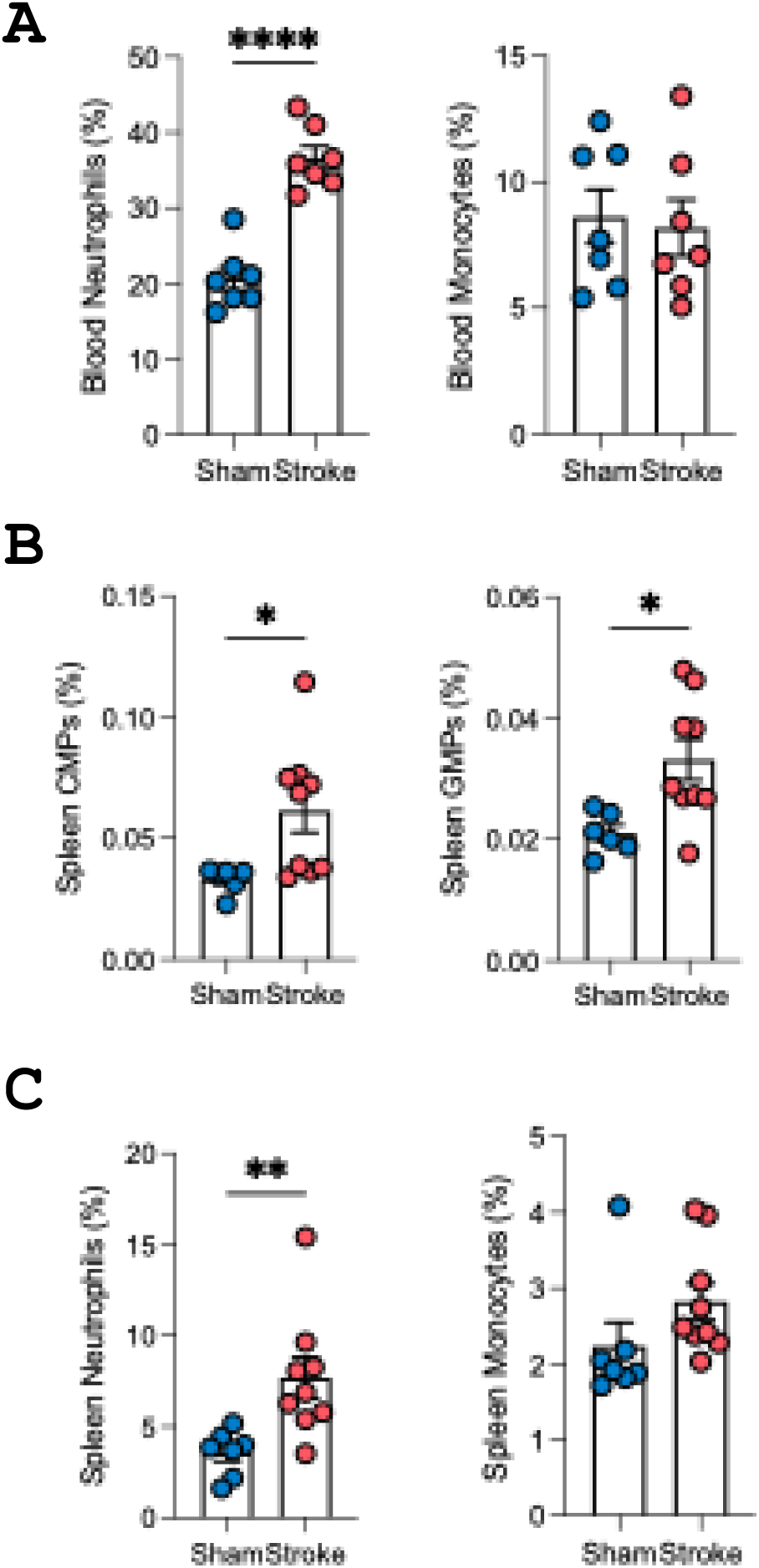
Stroke induces splenic, but not bone marrow, myelopoiesis. (A) Circulating leukocyte counts at 24 h show increased neutrophils, but no change in monocytes. (B) Splenic progenitors (myeloid progenitors, CMPs, and granulocyte–monocyte progenitors, GMPs) are significantly increased after stroke. (C) Splenic mature neutrophils also increased compared with sham, but no change in monocytes. Student’s t-test. *p<0.05, **p<0.01, ****p<0.001. n=6-9 per group.

### Inhibition of S100A8/A9 suppresses splenic myelopoiesis and systemic inflammation

Treatment with the S100A8/A9 inhibitor ABR-215757 markedly reduced splenic CMPs and GMPs (Figure 3A), as well as monocyte and neutrophil counts (Figure 3B), compared with vehicle-treated stroke mice. As expected, bone marrow myelopoiesis and mature myeloid cells remained unaffected (Supplementary Figure 2B). Inhibition of S100A8/A9 also significantly lowered circulating neutrophil and monocyte numbers (Figure 3C), confirming that blocking this pathway dampens systemic inflammation after stroke.

**Figure 3.**
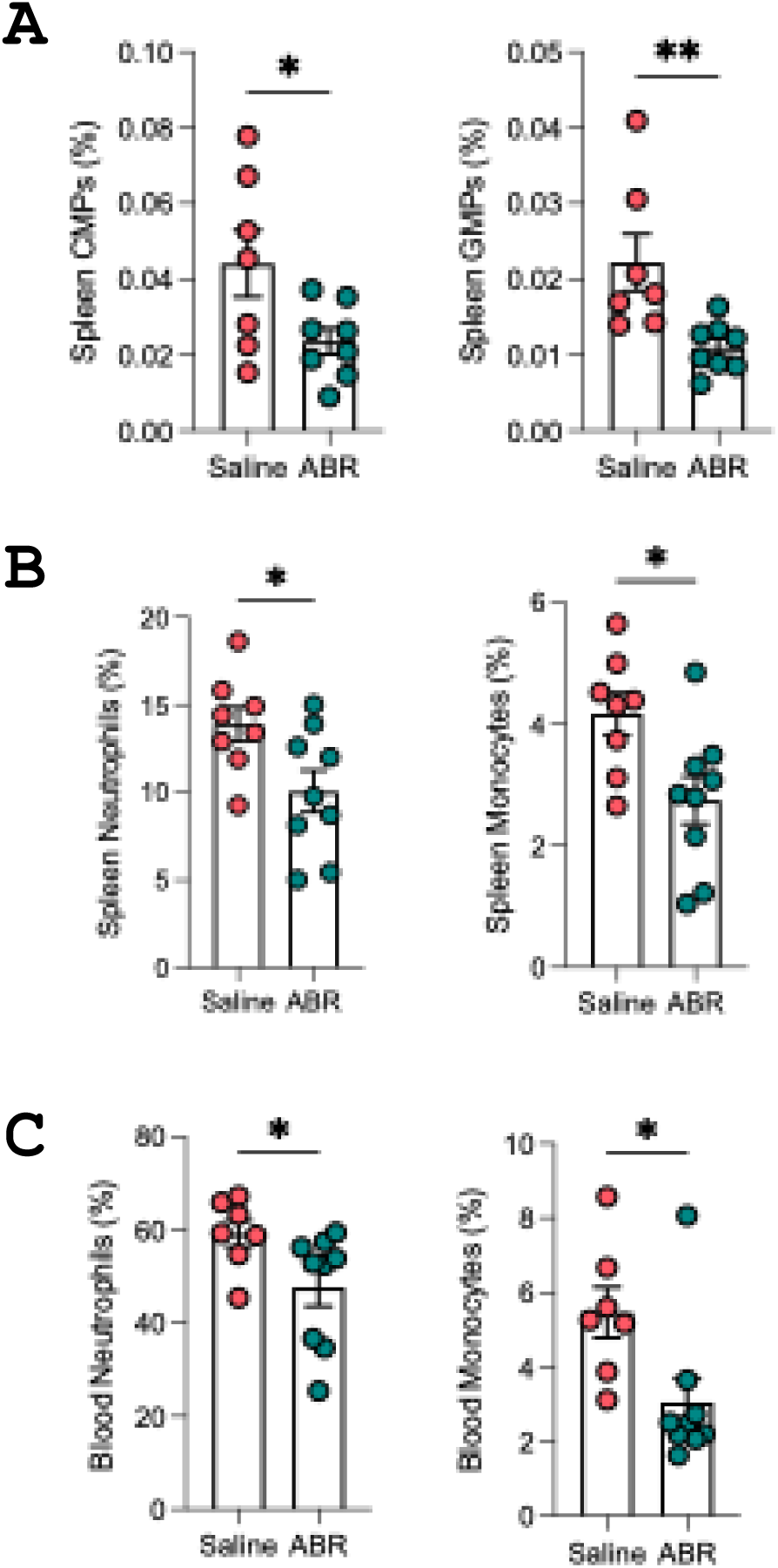
ABR-215757 reduces splenic myelopoiesis and systemic inflammation. (A) Splenic progenitors (CMPs, GMPs) are reduced by ABR-215757 treatment. (B) Splenic neutrophils and monocytes are significantly reduced by ABR-215757. (C) Circulating neutrophil and monocyte counts are also reduced by ABR-215757. Student’s t-test. *p<0.05, **p<0.01. n=7-9 per group.

### ABR-215757 improves stroke outcomes

Mice treated with ABR-215757 showed a ∼35% reduction in infarct volume compared with vehicle-treated controls (Figure 4 A and B), along with significantly improved neurological performance in the hanging wire test (Figure 4C). These findings demonstrate that inhibition of S100A8/A9 not only modulates peripheral immune responses but also provides functional neuroprotection in ischemic stroke.

**Figure 4.**
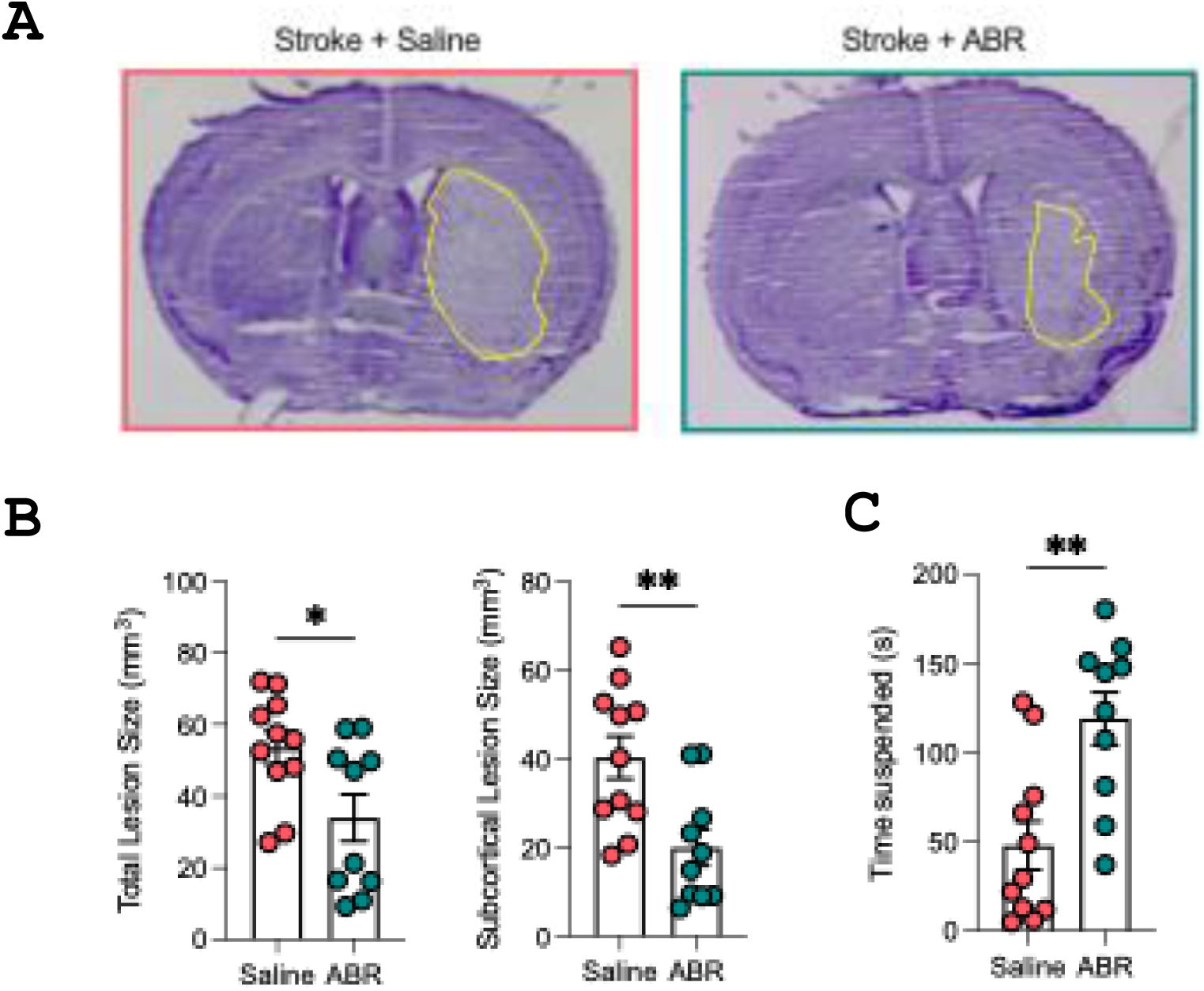
ABR-215757 improves outcomes after ischemic stroke. (A) Representative thionin-stained brain sections show reduced infarct size in ABR-215757–treated mice. (B) Quantification reveals ∼35% reduction in total and subcortical infarct volumes. (C) Hanging wire test shows significantly improved neurological function with ABR-215757. Student’s t-test. *p<0.05, **p<0.01. n=10-11 per group.

## Discussion

Ischemic stroke is a leading cause of disability, cognitive impairment, and mortality worldwide. Despite advances in our understanding of cerebral ischemia, current treatments focus almost exclusively on reperfusion strategies such as thrombolysis and thrombectomy, with no approved therapies targeting the immune response. Shortly after cerebral artery occlusion, ischemia triggers a cascade of inflammatory events, including endothelial activation, ROS production, and complement activation, which facilitate leukocyte recruitment and BBB disruption^26,27^. A growing body of evidence suggests that immune cells play complex and multiphasic roles, initially exacerbating tissue injury but later contributing to repair and regeneration^28^. Neutrophils are among the earliest immune cells to infiltrate the ischemic brain within minutes, peaking at 24-72 hours^29,30^, worsening injury through BBB disruption, oxidative stress, and release of proteolytic enzymes and their accumulation correlates with stroke severity and worse outcomes^31^. Consistent with this, we show an elevated number of neutrophils in the infarcted brain of both tMCAO mice and stroke patients, confirming that neutrophil-driven inflammation is a conserved feature of ischemic stroke. In this study, we demonstrate that S100A8/A9, a neutrophil-derived DAMP, is upregulated in the ischemic brain and promotes myelopoiesis in the spleen, leading to increased circulating neutrophils and monocytes. Importantly, we found that blocking S100A8/A9 with ABR-215757 suppressed splenic myelopoiesis, reduced circulating neutrophils, and significantly improved stroke outcomes. These findings provide novel insight into the regulation of stroke-induced immune responses and identify S100A8/A9 as a potential therapeutic target.

Beyond their role in brain injury, neutrophils contribute to systemic inflammation and myeloid expansion following stroke. We demonstrate that stroke leads to increased expression of S100A8/A9, a neutrophil-derived alarmin, within the infarcted brains of mice and humans. While we acknowledge that our human data on S100A8/A9 expression in the stroke tissue was limited to one sample, previous studies have shown a significant accumulation of S100A8/A9 in human stroke tissues^32^. Similarly, we confirmed elevated gene expression of *S100a8* and *S100a9* in the ischemic hemisphere in mice after tMCAO. S100A8 and S100A9 are primarily expressed in, and released from, myeloid cells upon cellular activation^15^. S100A9 gene expression is tightly regulated in a differentiation- and lineage-specific manner within the hematopoietic system^33^. Previously, we showed that induction of myocardial infarction promoted rapid recruitment of neutrophils and release of S100A8/A9 as alarmins. These complexes bind to TLR4 and prime the nod-like receptor family pyrin domain-containing 3 (NLRP3) inflammasome in neutrophils and promote IL-1β secretion^12^. The released IL-1β interacts with IL-1 receptor type 1 expressed on hematopoietic stem and progenitor cells in the bone marrow and stimulates granulopoiesis in a cell-autonomous manner. Our findings now extend this phenomenon to stroke, showing that S100A8/A9 is upregulated in the ischemic brain and likely functions as a systemic inflammatory signal driving myeloid expansion.

A central question in post-stroke immunity is the origin of circulating neutrophils. While emergency myelopoiesis has traditionally been attributed to the femoral bone marrow, it was shown that the skull bone marrow supplies neutrophils to the infarct within the first 24 hours^24^. Consistent with this, we observed no increase in CMPs, GMPs, or mature neutrophils in the femoral bone marrow 24 hours post-stroke. Instead, we found a striking expansion of CMPs, GMPs, and mature myeloid cells in the spleen, demonstrating that stroke triggers myelopoiesis as a contributing mechanism for increasing circulating neutrophils and monocytes. This aligns with previous reports that splenic contraction post-stroke releases stored leukocytes into circulation. Our findings also show that the spleen does not merely release pre-existing cells, but is an active site of myelopoiesis following stroke. While extramedullary myelopoiesis is a known phenomenon in chronic inflammation, such as in atherosclerosis and myocardial infarction, as demonstrated previously by our group^34,35^ and others^11,36^, its involvement in acute settings like stroke has not been previously reported. Our findings provide direct evidence that, in the absence of HSPC mobilisation from the bone marrow, the spleen can support the genesis of myeloid cells, beginning from CMPs.

After stroke, increased activation of the sympathetic nervous system (SNS) leads to reduced spleen size through splenic contraction and release of immune cells^37,38^. Other than this role of the SNS, our data suggest that S100A8/A9 could be a crucial factor that links brain ischemia to systemic myeloid expansion. Given its role in emergency myelopoiesis in myocardial infarction, it is likely that S100A8/A9 released from brain-infiltrating neutrophils acts as a systemic signal to promote myelopoiesis in the spleen. Therefore, we hypothesised that blocking S100A8/A9 signalling could suppress neutrophil production and reduce the severity of stroke outcomes. Using ABR-215757, an S100A8/A9 inhibitor that prevents the interaction with TLR-4 and RAGE^39^, we found that treatment significantly reduced CMP and GMP expansion in the spleen, reducing its eventual differentiation into circulating neutrophils and monocytes. Importantly, ABR-215757 treatment significantly reduced infarct volume development and improved neurological function. These findings highlight S100A8/A9 as a promising therapeutic target for reducing post-stroke inflammation and improving functional recovery.

Given the growing body of evidence linking neutrophils to infarct expansion and stroke severity, several strategies have been proposed to modulate neutrophil function as a therapeutic approach^40^. While direct neutrophil depletion has shown efficacy in preclinical stroke models, this approach carries risks of immunosuppression and increased infection susceptibility^41^. Our findings suggest that targeting upstream regulators of myelopoiesis, such as S100A8/A9, may provide a more selective approach to reducing harmful neutrophilia without compromising host defence. Beyond stroke, S100A8/A9 has been implicated in atherosclerosis, myocardial infarction, and thrombosis, suggesting that its inhibition could have broader implications for cardiovascular disease. Given that plasma S100A8/A9 levels correlate with infarct size and clinical outcomes in stroke patients, future studies should assess whether S100A8/A9 inhibitors could be translated into clinical trials for stroke and other vascular inflammatory disorders.

In conclusion, we identify S100A8/A9 as a key regulator of stroke-induced myelopoiesis, linking brain ischemia to systemic neutrophil production via extramedullary myelopoiesis in the spleen. We also demonstrate that blocking S100A8/A9 suppresses splenic myelopoiesis, reduces circulating neutrophils, and improves stroke outcomes. These findings further identify the spleen as a key organ in stroke and highlight S100A8/A9 as a novel therapeutic target to dampen post-stroke inflammation.

## Materials and Methods

### Human brain

Human brain tissue samples were obtained from an anonymized autopsy non-stroke subject and an anonymized acute ischemic stroke (AIS) patient receiving decompressive craniectomy and lobectomy at National Taiwan University Hospital with approval from the National Taiwan University Hospital ethics committee. The non-stroke subject was a 40 year-old male who had underlying severe aplastic anemia and died of peripheral blood stem cell transplantations with graft failure and multiple organ failure. The AIS patient was a 42 year-old male who had an infarction in the left middle cerebral artery (MCA) large territory and received surgical decompression and lobectomy two days after stroke. The brain tissues were collected within 48 h of death or at the time of surgery. Samples were taken from the temporal lobe within the infarcted area in the AIS patient. Brain tissues were fixed in 4% buffered formalin for at least three weeks before paraffin embedding. Brain sections were processed for immunohistochemical staining using primary antibodies against human CD15 [clone HI98] (Cat# 301902, Biolegend, US) and human S100A8/S100A9 heterodimer (Cat# MAB45701, R&D Systems, US) with a non-biotin-amplified method (Novocastra Laboratories Ltd, UK). Images were acquired using an Olympus microscope.

### Animals

This study fully adheres to the Animal Research: Reporting In Vivo Experiments (ARRIVE) guidelines^42^. All animal experiments were conducted in accordance with National Health and Medical Research Council of Australia guidelines for the care and use of animals in research and were approved by the Monash University Animal Ethics Committee. Mice had free access to water and food pellets before and after surgery.

### ABR-215757 treatments

ABR-215757 was a gift from Active Biotech and was used to therapeutically inhibit the bioactivity of S100A8/A9. Mice received drinking water containing ABR-215757 (0.5 mg/ml) or normal water 6 days prior to the induction of ischemia. ABR-215757 (0.5 mg/ml, 100 μl per dose) was injected intraperitoneally 24 h before and immediately after reperfusion. Control mice were injected with saline. Mice were randomized into different treatment groups and experiments were conducted in a blinded fashion.

### Transient focal cerebral ischemia

Focal cerebral ischemia was induced by transient intraluminal filament-induced occlusion of the right MCA, as described previously^43^. Mice were anesthetized with ketamine-xylazine (80 and 10 mg/kg, respectively; intraperitoneally). Rectal temperature was monitored and maintained at 37.5°C ± 0.5°C using an electronic temperature controller (Testronics, Kinglake, Victoria, Australia) linked to a heat lamp throughout the procedure and until animals regained consciousness. Briefly, the right proximal common carotid artery was clamped, and a 6-0 nylon monofilament with silicone-coated tip (Doccol Co., Redlands, CA, USA) was inserted and gently advanced into the distal internal carotid artery, 11-12 mm distal to the carotid bifurcation, occluding the MCA at the junction of the Circle of Willis. Severe (typically ∼80 %) reduction in rCBF was confirmed using transcranial laser-Doppler flowmetry (Perimed, Järfälla, Sweden) in the area of cerebral cortex supplied by the MCA. The filament was then tied in place and the clamp was removed. After 1 h of cerebral ischemia, the monofilament was retracted to allow reperfusion for 23h. Reperfusion was confirmed by an immediate increase in rCBF, which reached the pre-ischemic level within 5 min. The wound was then closed and the animal was allowed to recover. Regional CBF was recorded for 30 min of reperfusion. All animals were administered 1 mL of sterile saline via a subcutaneous injection for rehydration after surgery. Gel nectar (Able Scientific, WA, Australia) was placed inside the cage and access to chow food and water was provided. All animals’ boxes were placed on heat pads post-surgery until euthanasia. All mice were euthanized at 24 h by isoflurane overdose.

### Neurological assessment

At 23 h after induction of stroke, a hanging wire test was performed in which mice were suspended from a wire 30 cm high for up to 180 s, and the average time of 3 trials with 5-min rest periods in between was recorded. Neurological assessment was evaluated by an observer blinded to experimental groups.

### S100A8/9 and neutrophil staining

Mice were euthanized at 24 h by isoflurane overdose, followed by decapitation. The brains were immediately removed and snap frozen with liquid nitrogen. Coronal sections (10 μm) from vehicle-treated mice were fixed with 4% paraformaldehyde, endogenous peroxidase blocked, and stained using primary antibodies; rabbit recombinant multiclonal anti-S100A8+S100A9 [clone RM1038] (cat# ab288715, Abcam, UK) or rat monoclonal anti-mouse Ly-6G (cat# 551459, BD Biosciences, USA). Sections were then stained using horseradish peroxidase-conjugated secondary antibody against rabbit or rat (cat# P0448 or P0450, Dako, Denmark), respectively. Sections were incubated with a 3,3’-diaminobenzidine (DAB) substrate solution (cat# D4418, Sigma-Aldrich, USA) and counterstained with hematoxylin solution. Images were acquired using an Olympus light microscope.

### S100 gene expression

A separate cohort of mice were euthanized at 24 h by isoflurane overdose, and perfused with PBS via cardiac puncture. Ipsilateral (right) and contralateral (left) brain hemispheres were processed for RNA extraction. Total RNA was extracted using Trizol and cDNA synthesized using Superscript Vilo (Invitrogen, Thermo Fisher Scientific). Quantitative real time PCR was monitored in real time with an Mx3000 sequence detection system (Stratagene) using SYBR Green PCR Core Reagents (Agilent Technologies) and normalized to 18s.

*Flow cytometry*.

### Blood leukocyte counts and analysis

Neutrophils and monocytes were identified using flow cytometry as previously described^44^ . Blood was collected via tail bleeding into EDTA tubes, which were immediately incubated on ice. BM was harvested from the femurs and tibias. Spleens were flushed through a 40 μm cell strainer with PBS to obtain a single cell suspension^44^. For all organs, red blood cells were lysed using 1x RBC lysis buffer (BD pharm Lyse; BD Biosciences) for 15 minutes (blood) or 5 minutes (BM and spleen). Cells were then centrifuged at 400*g* for 5 minutes and resuspended in 1x HBSS containing 0.1% BSA w/v and 5 mM EDTA. Cells were stained with a cocktail of antibodies. For monocytes and neutrophil analysis, they were stained with anti-CD45 (PB), anti-Ly6-C/G (PerCP-Cy5.5) (BD Biosciences) and anti-CD115 (PE) (eBioscience). Monocytes were identified as CD45^hi^CD115^hi^, neutrophils were identified as CD45^hi^CD115^lo^Ly6-C/G^hi^ (Gr-1). For LSKs, CMPs and GMPs, a cocktail of antibodies to lineage committed cells (CD45R, CD19, CD11b, CD3e, TER-119, CD2, CD8, CD4, and Ly-6G; all FITC; eBioscience) and stem cell markers anti-Sca1 (Pacific Blue) and anti-ckit (APC-Cy7) were used. Haematopoietic stem and progenitor cells were identified as Lin^-^Sca1^+^ckit^+^ (LSKs). Where further identification of myeloid progenitor cells was required, antibodies to CD16/CD32 (FcγRII/III) and CD34 were used to separate common myeloid progenitors (CMPs) (lin^-^Sca1^-^ckit^+^FcγRII/III^int^CD34^+^) and granulocyte-macrophage progenitors (GMPs) (lin^-^Sca1^-^ ckit^+^FcγRII/III^hi^CD34^+^). All samples were run on a BD Canto II or BD LSR Fortessa X-20, and analysed using FlowJo (TreeStar). Gating strategy is shown in Supplementary Figure 3.

### Cerebral infarct and edema volume

Coronal sections (30 μm) separated by ∼420 μm were stained with thionin (0.1 %) to delineate the infarct. Images of the sections were captured with a CCD camera mounted above a light box. Infarct volume was quantified as described previously^45^ using image analysis software (ImageJ, NIH, Bethesda, MD, USA) and corrected for brain edema, estimated using the formula: corrected infarct volume = [left hemisphere area – (right hemisphere area – right hemisphere infarct area) × (thickness of section + distance between sections)]^45^. Edema-corrected infarct volumes of individual brain sections were added to give a three-dimensional approximation of the total infarct volume. Total and subcortical infarct volumes were quantified individually.

### Statistical analysis

Data are presented as mean ± SEM and were analyzed using the 2-tailed Student’s t test. P less than 0.05 was considered significant. All tests were performed using Prism software (GraphPad Software Inc.).

## Author contributions

H.A.K., A.A. and H.X.C. performed the experiments. H.A.K., A.J.M., C.G.S. and M.K.S.L. conceived the idea. H.A.K. and M.K.S.L. wrote the original draft of the paper. All authors reviewed and edited the paper.

## Acknowledgements

The authors were supported by Grants from the National Health and Medical Research Council of Australia (NHMRC).

---

**Supplementary Figure S1.**
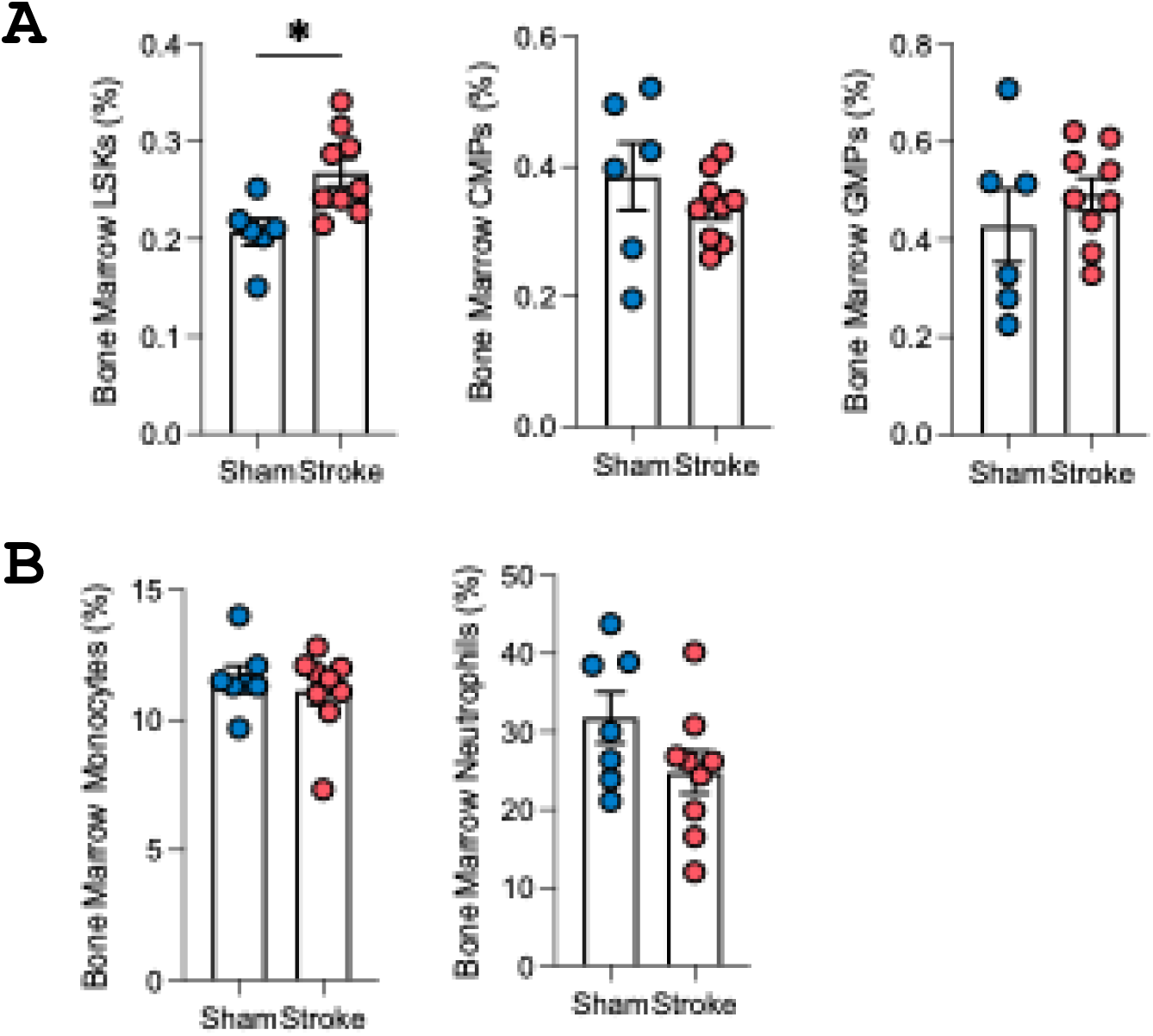

**Supplementary Figure S2.**
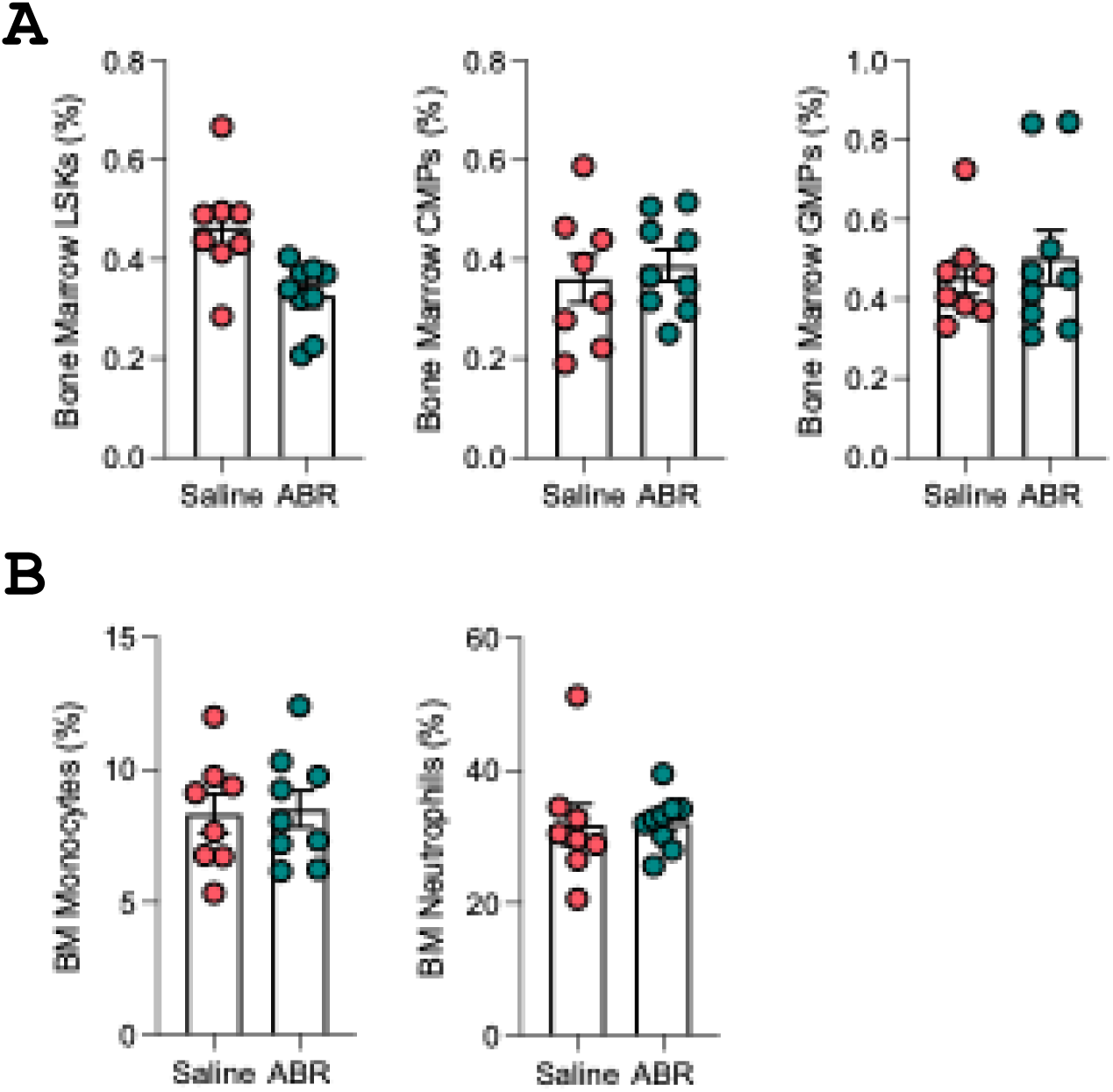

**Supplementary Figure S3.**
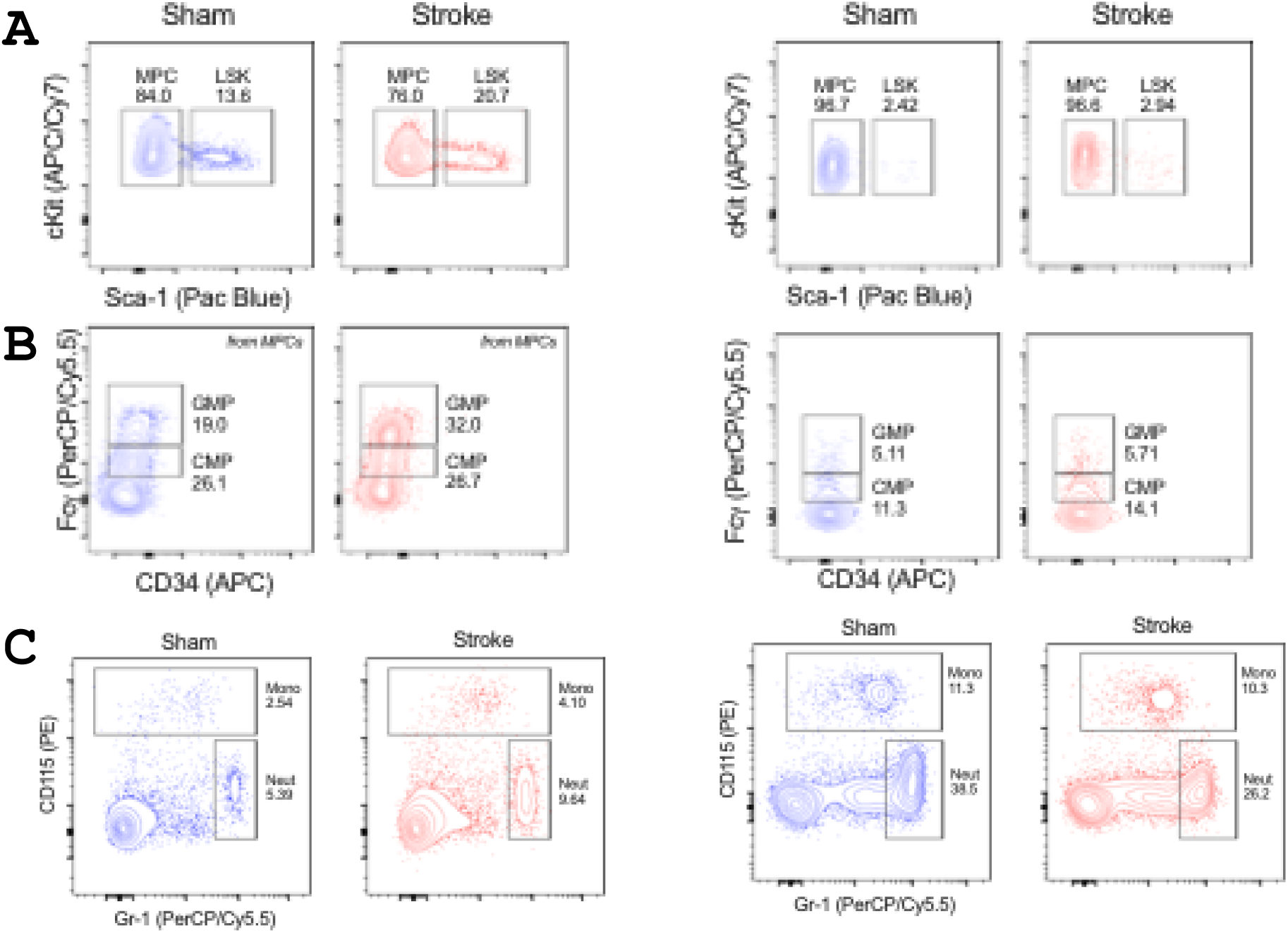

## Notes

### Competing Interest Statement

The authors have declared no competing interest.

